# Parental stress alters the fitness effects and genetic correlation of offspring performance traits and their gene expression pathways

**DOI:** 10.1101/2024.10.07.617098

**Authors:** Samuel N. Bogan, Marie E. Strader, Gretchen E. Hofmann

## Abstract

Transgenerational effects, where parental environments influence offspring phenotype, facilitate acclimation over ecological timescales. Transgenerational effects may also influence evolution by altering the fitness costs of offspring traits and the expression of genetic variation. We tested the evolutionary role of transgenerational effects in the sea urchin *Strongylocentrotus purpuratus*, whose populations are exposed to coastal upwelling (periods of low temperature and pH) and exhibit local adaptation, parental effects, and phenotypic plasticity in response to upwelling. Using a quantitative genetic breeding design, we conditioned parents and larvae to upwelling or control conditions and combined RNA-seq with larval phenotyping (body size, biomineralization, survival). Larval upwelling exposure caused widespread differential expression (DE), reduced biomineralization, and reduced size. Survival was linked to biomineralization, body size, and their plasticity under upwelling, but only in offspring of upwelling-conditioned parents -evidence that parental environment affected selection on offspring traits. DE under upwelling was associated with adaptive plasticity in biomineralization and size, but adaptive gene expression changes shared negative genetic correlations. However, genetic correlations in gene expression associated with body size plasticity were significantly more positive in larvae from upwelling parents. Parental conditioning strengthened selection on offspring performance and reduced genetic tradeoffs between performance-associated expression pathways, potentially accelerating adaptation.

## Introduction

The environmental experiences of parents can influence phenotypic outcomes in offspring through a process known as transgenerational plasticity (TGP). During TGP, the environmental cue triggering plasticity is experienced by a parent or parents and the plastic response to that cue is expressed in offspring and subsequent generations (Bell and Hellmann 2019). TGP has the potential to facilitate multigenerational, adaptive responses to environmental change over rapid ecological timescales, potentially aiding the resilience of biodiversity to global change (Donelson et al. 2018; Donelan et al. 2020). Additionally, TGP can influence natural selection, reshaping adaptation over evolutionary timescales (Day and Bonduriansky 2011; Herman and Sultan 2011; Donelson et al. 2018; Che et al. 2024). Theory regarding the evolutionary effects of parental environments has developed over three decades (Kirkpatrick and Lande 1989), but empirical research on the evolutionary consequences of transgenerational effects is lacking (Bogan and Yi 2024; Cao and Chen 2024). It is unclear whether and how strongly transgenerational effects alter the primary components of evolution by natural selection: the fitness effects and heritable genetic variation of traits.

Parental environments are predicted to reshape the fitness costs of offspring phenotypes. This can happen through alteration of parental investment in response to a stressful environment, with consequences for the condition and energy budget of offspring (Bonduriansky and Crean 2018). For example, offspring from parents that performed gametogenesis under oxidative stress may receive less maternal investment of lipids, proteins, and other resources (Schreck et al. 2001; Wong et al. 2019). If an offspring genotype underpins a phenotype that requires these resources, expression of that phenotype may come at a fitness cost (Buchanan et al. 2013). Assuming that there is heritable genetic variation in this phenotype, an effect of parental environment on the fitness effects of phenotypic variation will impact evolution by natural selection. For example, evidence that parental exposure to stress increases the fitness effects and expression of genetic variation in offspring traits would suggest that transgenerational effects can accelerate adaptation. However, an experiment has not been reported that evaluated the joint influence of transgenerational effects on the fitness costs and genetic variation of offspring traits.

Transgenerational effects can also alter the expression of genetic variation in offspring phenotype (Snell-Rood et al. 2016). An individual’s own environment is widely known to influence the expression of genetic variation. For example, environmental stress generally increases the expression of cryptic genetic variants (Rutherford and Lindquist 1998; Condelli et al. 2019). Therefore, genetic variation and covariation underpinning traits and their expression is dependent on the environment. Parental environments can alter the regulation of gene expression in offspring and phenotypes linked to that gene expression, resulting in the release or masking of genetic variation and changes in total additive genetic variation (Snell-Rood et al. 2016; Strader et al. 2022).

Changes in gene regulation that alter the expression of genetic variation have the potential to alter genetic covariance (Innocenti and Chenoweth 2013). Genetic covariance is a measure of how much two traits vary together as a function of genotypic effects. Under positive genetic covariance, genotypes that increase the expression of one trait generally increase the expression of a second trait. Under negative genetic covariance, genotypes exhibit opposing directional effects on two traits (Lande 1980). Genetic covariance is a critical constraint on evolution. Evolutionary rates accelerate when the directions of selection and genetic correlation of two traits align (e.g. positive directional selection on two traits sharing positive genetic correlation), but slow when the direction of these parameters is misaligned -e.g. positive directional selection on two traits sharing negative genetic correlation (Agrawal and Stinchcombe 2008; Blows and McGuigan 2016). If TGP induces differential expression in offspring of a gene with pleiotropic phenotypic effects, genetic covariance between those traits will change according to parental environment (Day and Bonduriansky 2011). If transgenerational effects alter the fitness consequences and genetic (co)variance of offspring phenotypes, then parental environments have potential to act as modulators of evolution by natural selection.

Performance traits and their phenotypic plasticity can be highly polygenic, regulated by multiple physiological pathways as identified by multi-environment genome-wide association studies (Li et al. 2018; Liu et al. 2021; Nguyen Ba et al. 2022) and gene expression-phenotype associations (DiLeo et al. 2011; Velotta et al. 2020). Whether a trait’s evolution is constrained or facilitated by genetic covariance depends, in part, on the strength and direction of covariance between the regulatory pathways underpinning it (Innocenti and Chenoweth 2013). RNA-seq experiments can be integrated with quantitative genetic breeding designs to (i) measure the additive genetic variation and heritability of regulatory pathways underpinning performance traits and (ii) estimate genetic variance-covariance (*G*) matrices of those pathways, consequently, their genetic correlation (Blows et al. 2015). Progress has been made in determining how developmental environments influence genetic correlation between traits (Sgrò and Hoffmann 2004; Wood and Brodie 2015) and their fitness effects (Kingsolver and Gomulkiewicz 2003). Whether parental environments reshape natural selection on traits through these mechanisms is poorly understood, however (Donelson et al. 2018).

Using the purple sea urchin *Strongylocentrotus purpuratus,* an ecologically important model invertebrate (Pearse 2006), we tested the hypotheses that parental exposure to stress reshapes the fitness effects and genetic (co)variance of offspring performance traits. We conditioned adults and larvae to experimental treatments that mimicked variation in temperature and *p*CO_2_ under coastal upwelling frequently experienced by *S. purpuratus*. Upwelling occurs when wind-driven, upward movement of deep seawater lowers the temperature and increases the *p*CO_2_ of surface oceans (Gruber et al. 2012). Coastal upwelling is predicted to increase in intensity as a result of climate change in regions such as the California Current (Snyder et al. 2003).

*S. purpuratus* populations inhabiting different *in situ* upwelling regimes exhibit genetically-fixed differences in their degree of phenotypic and transcriptional plasticity induced by experimental ocean acidification and upwelling, demonstrating that plastic, acclimatory response to this stressor are a key performance metric associated with upwelling adaptation (Evans et al. 2013, 2017; Kelly et al. 2013). In response to experimental upwelling, *S. purpuratus* exhibits transgenerational and developmental plasticity of differential gene expression, DNA methylation, and several performance traits including larval growth rate, biomineralization, and lipid content (Wong et al. 2018, 2019; Strader et al. 2019, 2020, 2022; Bogan et al. 2023). Phenotypic data for larval body size and biomineralization from the experiment that we describe here were reported by Strader et al. 2022, who detected significant genetic variation for the plasticity of both phenotypes in response to upwelling and an effect of parental environments on additive genetic variation in plasticity. Therefore, we evaluated our hypotheses by focusing on the plasticity of offspring phenotypes and gene expression as our primary performance traits.

We integrated RNA-seq with assays of performance and fitness-correlated traits in a quantitative genetic crossing design of larval families exposed to upwelling or control conditions (Figures 1A & S1), a design that facilitated the measurement of additive genetic variation, heritability, and *G* matrices associated with phenotypic plasticity and associated regulatory pathways (Fig. 1B). We tested whether parental stress modified selection on offspring plasticity and genetic covariance between pathways associated with offspring plasticity.

**Figure 1.**
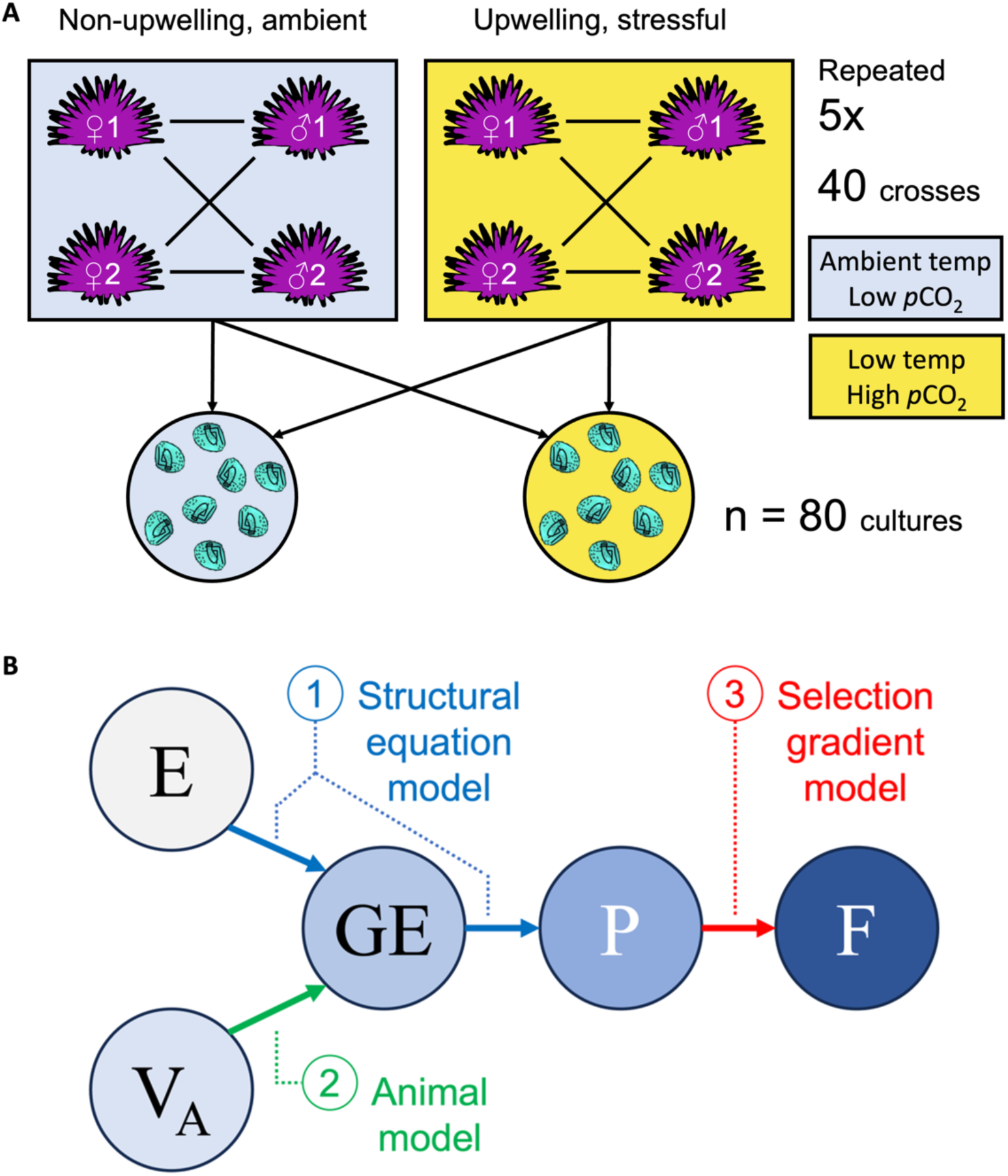
Crossing and experimental designs. **(A)** This graphic depicts adult and larval conditioning to ambient, non-upwelling (blue) and stressful, experimental upwelling conditions (yellow). Crosses between cohorts of conditioned adults are depicted with lines. Reciprocal rearing of offspring resulting from each cross is depicted with arrows directed toward larval non-upwelling and upwelling conditions. **(B)** A visual representation of modeling approaches used to integrate environmental treatments (“E”), additive genetic variance (“V_A_”), gene expression (“GE”), phenotypes (“P”), and fitness correlating traits (“F”). Structural equation modeling is visualized in blue which estimated the effect of differential expression on phenotypic plasticity induced by environmental treatments. Animal models predicting additive genetic variance in gene expression are green. Selection gradient models predicting the fitness effects of differential expression are red. Environmental treatments are grey. Parameters predicted by modeling approaches are visualized in shades of blue.

## 2. Methods

### 2.1 Adult conditioning, crossing, and larval culture

Adult urchins were collected from 2 coastal sites in the Santa Barbara Channel in August and September of 2018: Naples Reef (34.4221216, -119.95154) on August 23, 2018, and from Arroyo Quemado Reef (34.46774988, -120.11905). Adults were distributed across 4 90 L tanks per parental treatment at a density of 10 urchins per tank. Adults acclimated to parental treatments for approximately 4 months: non-upwelling = 17°C and 596 µatm *p*CO_2_; upwelling = 12.8°C and 1117 µatm *p*CO_2_. Flow rates to adult tanks equaled 20 l h^−1^. Adults were fed fresh *Macrocystis pyrifera ad libitum* with food changes and tank cleanings conducted once per week. Seawater temperature was controlled using heat pumps regulated by Nema 4X digital temperature controllers. *p*CO_2_ was controlled using a flow-through CO_2_ mixing system adapted from Fangue *et al*. (Fangue et al. 2010).

Fertilizations were conducted using a staggered cross-classified North Carolina II breeding design. During each phase of the staggered cross, 2 males and 2 females from a common adult condition were reciprocally crossed and their resulting offspring were cultured under non-upwelling and upwelling conditions until the prism stage of larval development. Each cross x larval treatment group was reared using 2 technical replicate culture buckets, resulting in 16 larval cultures per staggered cross. This process was repeated 5x for non-upwelling and upwelling crosses, alternating in order between parental treatments, resulting in 40 breeding adults, 40 crosses reared across 160 technical replicates, and 80 biological replicates (Fig. 1A). This sample size of breeding adults and lines exceeds breeding designs used to measure additive genetic variation and *G* matrices of transcriptome-wide gene expression in other animals (Blows et al. 2015; Tung et al. 2015).

Larvae were cultured in replicate flow-through 6 L nested buckets (i.e., an inner bucket with 30 µM mesh openings nested in an exterior bucket) at a flow rate of 3 L h^−1^ and a density of 10 larvae ml^−1^ until the early prism stage of larval development signified by the onset of tripartite gut differentiation. Temperature and pH were regulated in larval culture buckets as described above for adult conditioning. Point measures of temperature, salinity, total alkalinity, and pH for adult and larval tanks are described by Strader et al. 2022.

### 2.2. Phenotyping of performance and fitness-correlated traits

Three phenotypes were measured in larval cultures: (i) percent developmental abnormality (a corollary of survival), (ii) larval body size, and (iii) spicule length per unit body size (a corollary of biomineralization). Morphometric measurements of body size and spicule length were performed on *n* ≥ 30 prism larvae per technical replicated stored in 2% formalin buffered with 100 mM NaBO_3_ (pH 8.7) in filtered sea water. Body size was defined as the maximum linear distance of a prism body and spicule length defined as length from the tip of the body rod to the branching point of the post-oral rod. Abnormality was scored on *n* ≥ 30 larvae during sampling and was measured as the percentage of larvae exhibiting unsuccessful gastrulation. Gastrulation failure is frequently used as a proxy for larval viability and survival, as gastrulation is necessary for embryo viability and development (Duncan et al. 1997; Gambardella et al. 2021). Importantly, body size and spicule length were not measured in non-prism larvae or ungastrulated embryos. Because RNA-seq was performed using pooled RNA samples per culture, performance and fitness-correlating phenotypes were integrated with gene expression data using culture means rather than per-animal values.

### 2.3. RNA extraction, sequencing, and analysis

Total RNA was extracted with Trizol from pooled samples of 6,000 larvae per culture replicate. Extractions were performed on 1 technical replicate per cross x developmental treatment resulting in 80 RNA extractions. Total RNA quantity and quality was evaluated via Nanodrop, gel electrophoresis, and Qubit quantification before library preparation. RNA-seq libraries were prepared using polyA enrichment and were quality checked via LabChip GX. Strand specific PE 150 bp reads were sequenced on an Illumina HiSeq 4000 platform. Adapter removal, quality filtering, and trimming were performed using CutAdapt v4.4 and Trimmomatic v0.39 before reads were aligned to the ‘Spur_5.1’ reference genome (Sodergren et al. 2006) using hisat2 v2.2.1 (Kim et al. 2019). Specific parameters for read processing, alignment, and processing of alignment files are reported in Supplemental Methods.

Transcript read counts were normalized in edgeR v3.40.2 as counts per million (CPM). Read counts were filtered to keep all transcripts exhibiting CPM > 0.5 in at least 75% of the 80 replicates. Differential expression (DE) was modeled using a negative binomial generalized linear model (glm) fitted with edgeR’s robust, tagwise dispersion parameter using the robust iteration of the model fitting function ‘GLMQLFit’ and the DE test function ‘GLMQLFTest’ (Robinson et al. 2010). Expression was predicted as a function of two non-interacting, categorical variables for parental and developmental environment. Models were fit with non-interacting environmental predictors because the study’s design only enabled the measurement of V_A_ for developmental rather than transgenerational plasticity. Fitting an interaction between both effects would confound interpretation of V_A_ for developmental plasticity. Significant DE was evaluated using FDR adjusted p-values (alpha < 0.05). Functional enrichment was tested using a rank-based Mann Whitney U test of Gene Ontology terms input with logFC coefficients for DE. This test determines whether a given GO term’s logFC distribution is significantly skewed from the mean of the background, filtered transcriptome (Wright et al. 2015).

### 2.4 Identifying adaptive differential expression

The effect of DE on the plasticity of body size and biomineralization (body size-normalized spicule length) was measured using structural equation models (SEMs). SEMs were derived from two linear models: (i) phenotype predicted as a function of transcript abundance, developmental environment, and parental environment and (ii) scaled, signed transcript abundance predicted as a function of developmental and parental environments. Scaled transcript abundance was signed such that samples with low expression resulting in a negative scaled value were multiplied by the direction of the transcript’s DE under upwelling. The effect of developmental environment on phenotype mediated by DE was estimated for each transcript using mediation analysis performed with the ‘mediate’ function of the R package mediation v4.5.0 set to 1000 simulations (Tingley et al. 2014). Positive mediation effects indicated that changes in gene expression in the direction of DE were associated with higher levels of body size or biomineralization under upwelling. Negative mediation effects indicated that DE was associated with reduced phenotypic values under upwelling. To understand how the strength of DE impacted phenotypic outcomes, linear regressions were performed between transcriptome wide, absolute logFC and a second-order, quadratic polynomial for the phenotypic effects output by SEM.

The fitness costs and benefits of plasticity in body size and biomineralization were measured using an adaptation of the Lande & Arnold selection gradient model (Lande and Arnold 1983), which traditionally assumes uniform relatedness among individuals, for use on non-uniformly related samples. This was achieved by controlling for the effect of genetic covariance on the fitness-correlating outcome using a known pedigree structure. A fitness correlated trait (proportion of normal development, a larval corollary of survival) was modeled as a function of developmental and parental environment, body size or biomineralization per cross in each environment, and the plasticity of body size or biomineralization of a cross between developmental environments. Selection gradient models included a random effect identifying each cross and controlling for genetic covariance using a relatedness matrix generated from custom code (see https://github.com/[REDACTED]) such that offspring-parent relatedness equals 0.5, full-sibling relatedness equals 0.5, half sibling relatedness equals 0.25, and unrelated cohorts share a relatedness of 0. This modification of the Lande and Arnold model has been used to accommodate non-uniform relatedness in other studies (Fontúrbel et al. 2021).

Selection gradient models were fitted in brms v2.19.6 (Bürkner 2017), an R interface to the Bayesian programming language Stan (Carpenter et al. 2017). Models assumed uniform priors, employed 40,000 MCMC iterations with a 5,000 iteration warm up, and a beta-distributed generalized linear regression model family. Beta distribution was selected because the proportion of normal development is constrained between 0 and 1. A Bayesian approach was selected for fitting because of the flexibility of packages such as brms for accommodating relatedness matrices within a beta distributed model family. These selection gradient models predicted whether positive versus negative plasticity of larval body size and biomineralization promoted greater fitness under upwelling stress. The significance of fitness effects were tested using probability of direction, which determines whether the ≥ 95% of posterior distribution falls above or below 0 (Makowski et al. 2019). Selection gradient coefficients were then multiplied with the phenotypic effects of DE on plasticity for body size and biomineralization to calculate the associated fitness effect of transcriptional plasticity.

### 2.5 Measuring the heritability of gene expression and its plasticity

Additive genetic variation (V_A_) and heritability (*h*_2_) of gene expression was measured to determine whether there was sufficient genetic variation for fitness-and phenotype-associated changes in expression to evolve via natural selection. V_A_ for gene expression and DE were measured across all transcripts using animal models fit with the ‘relmatLmer’ function of the R package lme4qtl v0.2.2 (Ziyatdinov et al. 2018). Within animal models, DE was measured as the foldchange of gene expression across developmental environment for a given cross. Mean-standardized CPM (gene expression) was predicted as a function of fixed effects for developmental and parental environment, random effects for dam and sire, and a random effect for cross identity. Genetic covariance between crosses was estimated using the relatedness matrix described above. Mean-standardized DE fold changes were predicted using an identical animal model lacking a fixed effect for developmental environment. *h*^2^ was derived from each model as the heritable proportion of total variance in gene expression or DE. Standard error in *h*^2^ was calculated for each model via bootstrapping using the function bootMer function of lme4set to “parametric” and 100 simulations (Bates et al. 2015). Differences in *h*^2^ of baseline gene expression and gene expression were modeled transcriptome wide using a quasibinomial glm linear model that corrected for standard error of *h*^2^ by weighting observations by 1/SE^2^.

Covariation between DE’s heritability and adaptive, phenotypic effects were modelled using three different approaches addressing the questions (i) ‘Does the probability of heritability (*h*^2^ ý 0.2) vary as a function of DE’s effect on adaptive plasticity in body size or biomineralization?’, (ii) ‘Does total heritability (*h*^2^ as a continuous variable) vary as a function of DE’s phenotypic effects?’, and (iii) ‘Does total heritability vary according to the fitness costs of DE’s combined effect on plasticity in body size and biomineralization?’. Tests of questions i and ii were performed by modelling the probability of *h*^2^ ≥ 0.2 using a binomial glm. Continuous *h*^2^ was modeled as a quasibinomial linear model weighted by *h*^2^ standard error (weight = 1/SE^2^). Each model type fitted two continuous predictor variables for the phenotypic effect of DE on body size and biomineralization signed toward the adaptive direction of that effect under parental upwelling, which induced yielded fitness effects of plasticity as opposed to neutral effects under parental non-upwelling. Models of continuous *h*^2^ set parameters for DE’s phenotypic effects as a second order polynomial to accommodate non-linear variation in *h*^2^ across the parameter space. For question iii, DE’s total effect on adaptive plasticity was calculated as the summation of the SEM-predicted coefficient for DE of transcript *i*’s effect (*E*) on body size (*S*) and biomineralization (*B*) multiplied by the selection gradient (*β*) acting on each plastic trait under upwelling, such that adaptive plasticity associated with transcript *i* = (*E_S,i_* x *β_S,i_*) + (*E_B,i_* x *β_B,i_*). Covariance between DE *h*^2^ and its total adaptive effect on plasticity was modeled using a quasibinomial linear model weighted by standard error of *h*^2^ (weight = 1/SE^2^). The predictor variable of this model was calculated as the sum of two products that were separately measured for the plasticity of biomineralization and size. For each trait, this product equaled the indirect effect of DE on plasticity multiplied by the selection gradient acting on plasticity. For example, this predictor variable was high when DE had a strong and positive association with plastic decreases in upwelling, which were adaptive.

### 2.6. Estimating G matrices

Genetic variance-covariance matrices of absolute logFC values for DE associated with adaptive plasticity in larval body size and biomineralization were estimated by fitting multivariate animal models representing all possible pairs of DEGs in each group. Additive genetic variance in each absolute logFC trait and covariances between pairs were extracted from the multivariate models and assembled into *n*-by-*n* variance-covariance matrices, where *n* equals the number of DEGs associated with adaptive plasticity in body size or biomineralization. Genetic variance-covariance matrices were then converted to correlation matrices using the cov2corr() function of R ‘stats’ v4.2.2. The number of genetically-correlated modules of adaptive plasticity-associated DEGs was estimated from *G* matrices using hierarchical clustering of genetic distances (1 – genetic correlation). A distance threshold of < 0.5 was used to assign DEGs to their shared modules. Hierarchical clustering was performed using the hclust() function of R stats v4.2.2.

Variation in logFC *G* matrices between larvae of upwelling and non-upwelling parents was quantified. First, two multivariate animal models fitted with pedigrees of parental non-upwelling or parental upwelling families were fitted for all pairs of genes with DE associated with adaptive plasticity in larval body size or biomineralization. As described above, genetic (co)variances from each model were used to populate parental upwelling and non-upwelling *G* matrices which were converted to correlation matrices. Significant differences in correlation matrix structure between parental treatments were evaluated using a three-step approach. Two-sided Mantel tests were used to compare parental environment matrices, applying separate tests to logFC associated with adaptive plasticity in body size and adaptive biomineralization plasticity. Mantel tests were run using the mantel.test() function of the R package ‘ape’ v5.8 using 10,000 iterations (Paradis et al. 2004; Paradis and Schliep 2019). Next, a chi-squared test was used to determine if the number of negative-to-positive changes in genetic correlation under upwelling relative to positive-to-negative changes was greater than expected by chance.

To determine whether significant enrichment in negative-to-positive changes in genetic correlation was attributed to parental environment or the pedigree structure of populations in parental treatment groups (i.e., differences in genetic (co)variance between animals used in each treatment), random permutations of the experimental pedigree were constructed. These permutations split the pedigree into two halves which evenly represented families of parental treatment groups. Random permutations of split pedigrees were input into animal models and resulting *G* matrices. Chi-squared tests of enrichment for directional changes in genetic correlation between permuted matrices were performed, which determined the effect of pedigree structure on apparent effects of parental environment on genetic correlations. In Results, enrichment of directional changes in genetic correlation between parental environment are reported alongside the percentage of these effects that were attributed to pedigree structure rather than treatment.

## 3. Results

Coastal upwelling is stressful for *S. purpuratus* larvae (Wong et al. 2018, 2019; Strader et al. 2020, 2022). This was similarly observed here, as evidenced by an increased percent abnormality in larvae in cultures conditioned to the treatment transgenerationally and developmentally (Strader et al. 2022). Developmental and parental conditioning to experimental upwelling induced widespread differential expression (DE; Fig. 2). DE induced by developmental upwelling was primarily associated with decreases in larval body size and, to a lesser extent, increases in biomineralization (Fig. 3). The plasticity of body size and biomineralization both yielded significant fitness effects measured as variance in the proportion of normal development among larval families. However, fitness effects were contingent upon the parental environment from which larvae were spawned. Following parental upwelling, reductions in larval body size and plastic increases in biomineralization were both adaptive. Using a heritability threshold of *h*^2^ ý0.2 (Orton 2020), 56.80 – 59.93% of adaptive gene expression plasticity exhibited significant heritability (Fig. 4). Negative genetic correlation was abundant between DE of genes associated with adaptive phenotypic plasticity. However, larvae from upwelling-exposed parents exhibited a significant reduction in negative genetic correlation between DE genes associated with adaptive plasticity in body size, demonstrating that changes in gene regulation following parental stress modified the expression of genetic covariance (Fig. 5). We further describe these results below.

**Figure 2.**
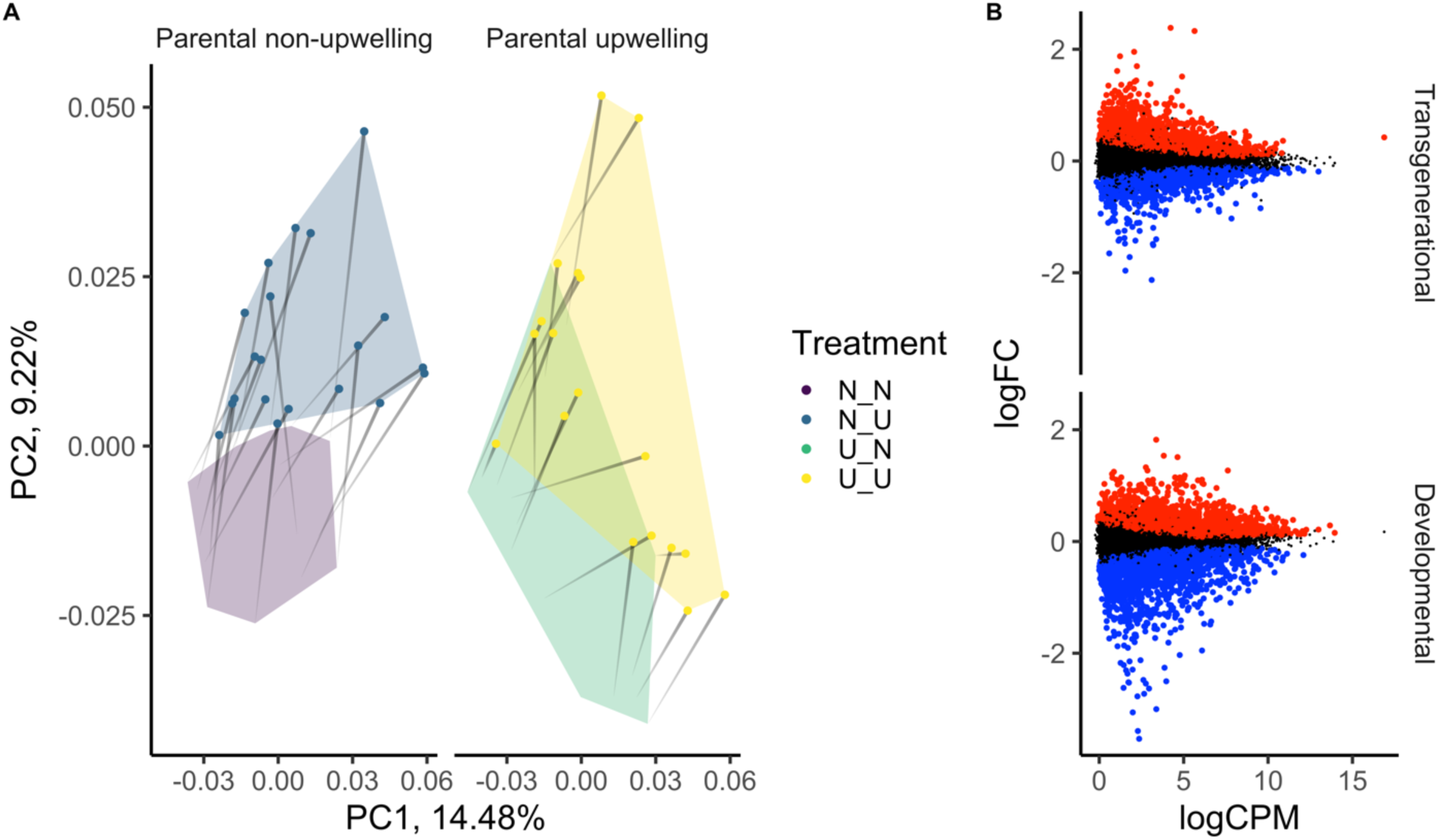
Differential expression induced by parental and developmental exposure to upwelling. **(A)** Loading of samples to principal coordinate axes derived from filtered, normalized read counts. Parental and developmental treatments are represented by color. Paths connect single crosses and their change in loading between non-upwelling (“N”; no point) and upwelling (“U”; point) treatments. **(B)** Mean difference plots of transcript logFC across average CPM per transcript, grouped by parental and developmental effects of upwelling. Significant downregulation is depicted in blue and upregulation in red.

**Figure 3.**
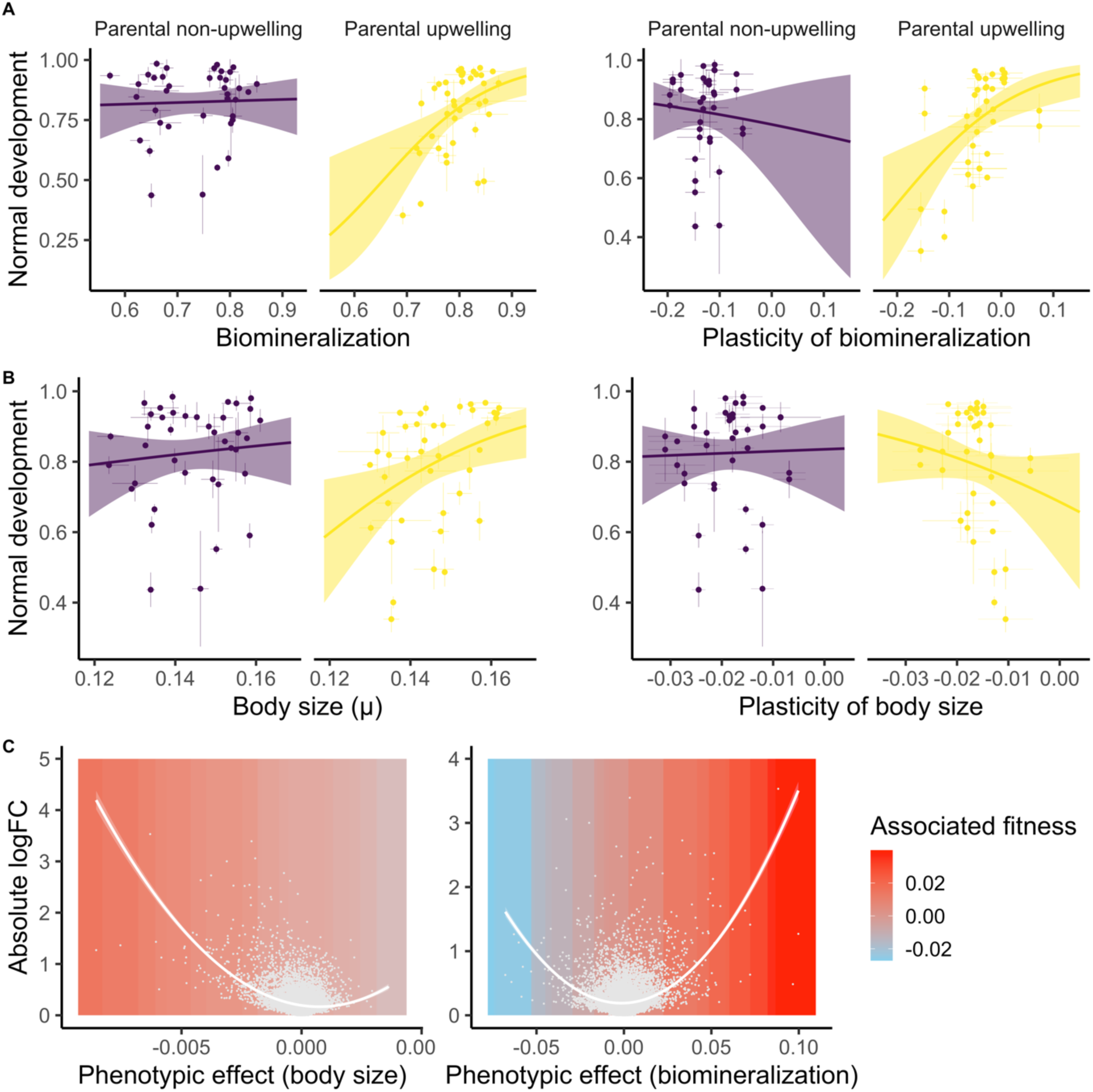
Effects of differential expression on adaptive plasticity. **(A – B)** The effects of phenotypic values and plasticity in biomineralization (spicule length normalized by body length) and body size (maximum body length) on proportion of normal development grouped by parental environment are plotted in A and B, respectively. Points represent phenotypic means (left) or mean reaction norms (right) of crosses reared in each environment. Environments are depicted by color such that non-upwelling is blue and upwelling is yellow. Error bars depict standard deviation in each trait among replicates of each cross. **(C)** Absolute logFC of differential expression induced by developmental upwelling is plotted across differential expression’s association with upwelling effects on body size and biomineralization in A and B, respectively. Points represent transcripts. Solid lines depict fitted quadratic curves ± 95% CI. Plot background color corresponds to the product of differential expression’s phenotypic effect on the plasticity of body size or biomineralization and the selection gradient for plasticity of each trait under parental upwelling, resulting in an inferred fitness induced by differential expression.

**Figure 4.**
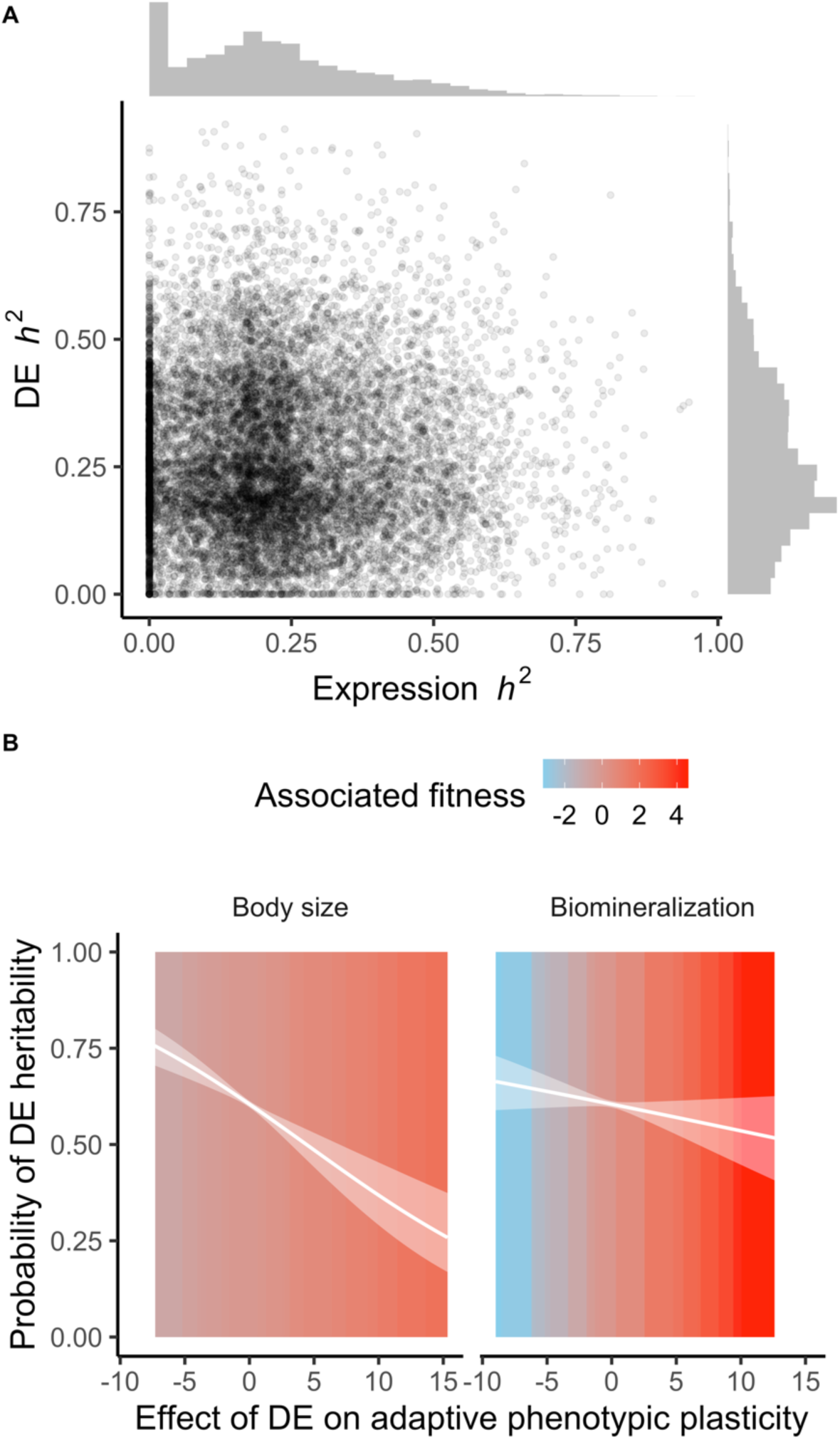
Heritability of differential expression and its association with adaptive phenotypic plasticity. **(A)** *h*^2^ of differential expression (DE) induced by developmental upwelling is plotted across *h*^2^ of baseline gene expression transcriptome wide. The distributions of *h*^2^ for baseline and differential expression are plotted adjacent to the x and y axes, respectively. **(B)** The probability of differential expression *h*^2^ ≥ 0.20 is plotted as a logistic curve ± 95% CI across differential expression’s effect on biomineralization and body size signed for the effect’s direction toward adaptive plasticity. Background color represents the coefficient for DE’s effect on phenotypic plasticity multiplied by the selection gradient acting on the plasticity of each phenotype under parental upwelling (e.g., the associated fitness outcome of DE’s effect on plasticity).

**Figure 5.**
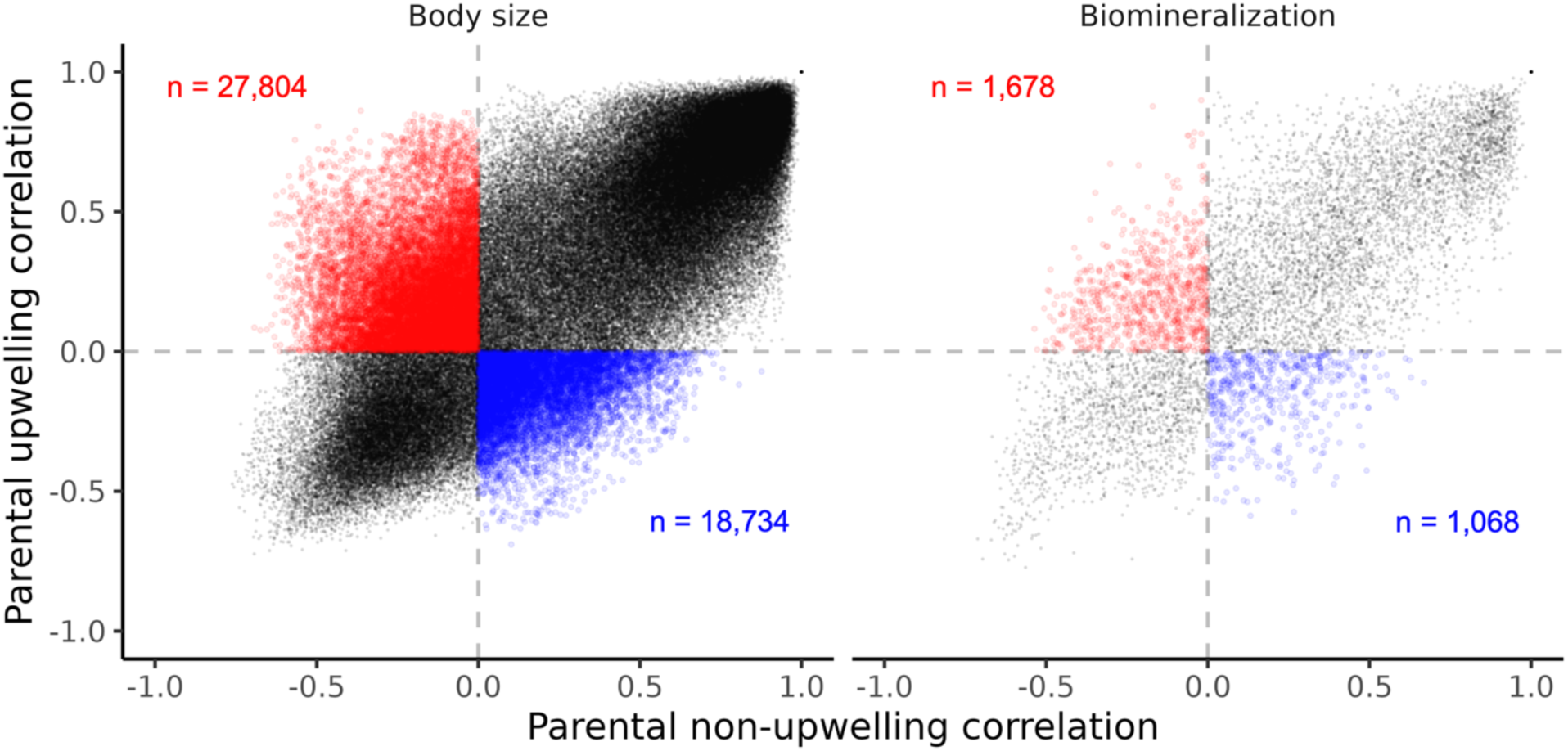
Changes in genetic correlation between parental upwelling treatments for log foldchanges (logFC) of differential expression (DE) associated with adaptive plasticity in larval body size and biomineralization. X and Y axes represent genetic correlations extracted from *G* matrices of logFC values estimated from larvae of non-upwelling and upwelling conditioned parents. Correlations that shifted from negative to positive under parental upwelling are enlarged and colored in red. Correlations that shifted from positive to negative under parental upwelling are enlarged and colored in blue.

### 3.1 Differential expression induced by parental and developmental upwelling

*RNA-seq alignment, quality checking, and filtering* – Following trimming, RNA-seq libraries achieved a mean size of 37.07 ± 5.09 million reads and a mean mapping efficiency of 74.14 ± 1.80%. After read-count filtering for transcripts with > 0.5 CPM in at least 75% of all samples, 12,953 transcripts were retained for downstream analysis. Outlier detection was performed with filtered read count data using arrayQualityMetrics v3.54.0 (Kauffmann et al. 2009), which flagged two half sibling crosses from parental and developmental upwelling treatments as significant outliers (Fig. S2–S3). Removal of these two samples brought the RNA-seq sample size to n = 78. Mean variation in gene expression across samples equaled a biological coefficient of variation of 0.12. Library and alignment metrics are further described in Supplemental File 1.

Parental conditioning to upwelling induced upregulation of 1,582 transcripts and downregulation of 1,539 in larval offspring. These differentially expressed genes (DEGs) included 50 and 29 upregulated and downregulated transcripts with an absolute log_2_FC > 1.0 (Fig. 2). DEGs induced by parental upwelling were enriched for 71 biological process (BP) GO terms, 49 molecular function (MF) terms, and 30 cellular component (CC) terms described in Figures S4– S6. Developmental exposure to upwelling induced upregulation of 2,246 transcript and downregulation of 2,205. With a >1.0 log_2_FC cutoff, these included 38 upregulated and 184 downregulated transcripts (Fig. 2). In comparison of parental and offspring responses to upwelling, 39.73% of DEGs induced by transgenerational effects were also differentially expressed in response to developmental conditioning. Developmental upwelling DEGs were enriched for 89 BP GO terms, 53 MF terms, and 37 CC terms described in Figures S7–S9.

Upregulated transcripts related to ribosomal function included several ribosomal subunits and 16 DEAD-box proteins involved in the initiation of translation. Included in cellular responses to stress was the significant upregulation of 8 heat shock proteins including 3 Hsp70 and 5 Hsp40 chaperones. Interestingly, 8 heat shock proteins were significantly downregulated in response to upwelling, indicating that they were more highly expressed under warmer conditions. These included all 3 Hsp90 isoforms present in the *S. purpuratus* genome as well as 2 Hsp70 and 3 Hsp40 chaperones. These functional enrichment results demonstrate a suite of complex molecular responses to multivariate, abiotic environmental change brought on by experimental upwelling, a third of which were commonly induced by parental and larval conditioning.

### 3.2. Selection on offspring performance traits and their plasticity

Selection gradient models predicting the proportion of normal development per replicate culture (a fitness correlated trait) as a function of body size or biomineralization and their plasticity estimated fitness benefits of (i) plastic reductions in size under upwelling and (ii) maintenance of or plastic increases in biomineralization under upwelling. When larvae were spawned from parents exposed to upwelling, larval biomineralization incurred a selection gradient of 0.15 ± 0.06 (Fig. 3A) and, for body size, 0.13 ± 0.04 (Fig. 3B). In larvae from upwelling parents, plastic increases in biomineralization incurred a selection gradient of 0.40 ± 0.09 (Fig. 3A) and plastic reductions in body size incurred a weaker selection gradient of 0.13 ± 0.04 (Fig. 3B). Interestingly, the plasticity of both traits did not exhibit detectable fitness costs or benefits when larvae were spawned from non-upwelling parents. These selection gradient coefficients represent the slopes of logistic regressions between larval survival (abnormality) and phenotypic plasticity visualized in Figures 3A and 3B.

### 3.3. Associations between differential expression and adaptive versus maladaptive plasticity

Structural equation models (SEMs) predicting the effect of differential expression (DE) on plastic changes in body size and biomineralization in response to developmental upwelling identified 231 transcripts associated with increased body size, 564 with reduced body size, 125 associated with increased biomineralization, and 113 with decreased biomineralization. Because selection gradients were significantly positive for plasticity of biomineralization, and negative for plasticity in size, DE associated with reductions in body size or increases in biomineralization were associated with adaptive or weakly adaptive phenotypic outcomes following parental exposure to upwelling (Fig. 3C).

As the absolute fold change of a transcript’s DE increased, the adaptive effect of DE on reduced body size and/or increased biomineralization significantly increased. From here forward, ‘adaptive’ or ‘maladaptive’ effects refer to selection on larvae from upwelling-exposed parents unless otherwise specified. Associations between DE and effects on phenotypic plasticity toward maladaptive directions were weaker, demonstrating that DE induced by developmental upwelling was predominantly associated with adaptive plasticity (Fig. 3C). Transcripts associated with reduced body size under upwelling conditions were enriched for BP/MF terms involved in cellular signaling, transmembrane transport, and ribosomal biogenesis localized to cell junctions and the nucleolus. DE driving increases or decreases in biomineralization exhibited no functional enrichment across all GO term categories.

### 3.4. Genetic (co)variance of adaptive and maladaptive differential expression

DE was highly heritable across most genes, but levels of heritability varied according to associations between DE and adaptive plasticity. DE was more heritable than baseline gene expression (CPM) by 19.95% *h*^2^ (p < 2.2e^-16^) and the two variables were uncorrelated (Fig. 4A). Most significant DE induced by developmental conditioning (60.46%) exhibited *h*^2^ greater than or equal to 0.20 (i.e., at least 20% of in DE variance was heritable). The mean *h*^2^ of expression was 0.2229 ± 0.1159. The standard error of expression *h*^2^ was 0.1159 ± 0.0486. The mean *h*^2^ of DE was 0.2674 ± 0.1592. The standard error of DE *h*^2^ was 0.1158 ± 0.0486. 7.59% of DEGs were heritable and associated with adaptive plasticity in body size. However, 59.93% of adaptive DE related to body size was heritable. These transcripts exhibited similar *h*^2^ relative to all DE: 0.2613 ± 0.1516. Less DE was heritable and associated with maladaptive plasticity in body size -only 3.59% of all DEGs. However, maladaptive DE related to body size exhibited greater heritability with a mean *h*^2^ of 0.2984 ± 0.1600. Most of the DEGs associated with adaptive increases in biomineralization (56.80%) were heritable, with a mean *h*^2^ of 0.3709 ± 0.1224. 1.55% of DEG’s were heritable and associated with adaptive increases in biomineralization. No DEGs that were significantly associated with maladaptive decreases in biomineralization were heritable.

DE associated with adaptive reductions in body size under upwelling was less heritable than DE associated with maladaptive increases in size. As the effect of DE on adaptive decreases in body size grew, *h*^2^ significantly decreased (*β* = -0.96; p = 5.77e^-8^) as well as the probability of *h*^2^ *β* 0.2 (*β* = -5.47; p = 2.16e^-8^; Fig. 4B). The phenotypic effect of DE on adaptive biomineralization plasticity had a negative but insignificant effect on its heritability (Fig. 4B). Adaptive DE remained significantly less heritable when considering the cumulative fitness effects of plasticity-associated DE (*β* = -0.05; *p* = 0.00376). The cumulative fitness effect of DE was measured by summing DE’s phenotypic effects on both traits and multiplying these effects by the selection gradients acting on each trait’s plasticity.

Genetic variance-covariance matrices (*G* matrices) were estimated for sibships’ logFC values for differentially expressed genes associated with adaptive plasticity of body size and biomineralization. DE associated with adaptive plasticity in body size was composed of 28 genetically correlated modules of 564 genes. DE associated with adaptive plasticity in biomineralization was composed of 32 genetically correlated modules of 125 genes. Genetic correlations between logFC values were primarily positive and exhibited larger absolute correlation among logFCs sharing positive genetic correlation (Fig. 5). *G* matrices for larvae from upwelling versus non-upwelling parents exhibited differences in genetic correlation for DE associated with adaptive plasticity in body size. Mantel tests were used to evaluate variation between parental treatments in genetic correlation matrices of DE associated with adaptive plasticity in body size (*Z* = 40477.59; *p* < 1e10^-4^) and biomineralization (*Z* = 1438.176; *p* < 1e10^-4^). For both traits, some correlations shifted from negative-to-positive and positive-to-negative under parental upwelling (Fig. 5). Even in the absence of environmental effects, random chance in changes to the sign of genetic correlation are expected. Thus, a chi-squared test was used to determine if there were greater or fewer positive-to-negative shifts than expected by chance alone. Genetic correlations in body size DE exhibited significant shifts from negative-to-positive under parental upwelling (χ^2^ = 1767.7; *p* < 2e10^-16^).

Parental treatment groups contained different genotypes, potentially confounding transgenerational effects on genetic correlation. Permutations of the experimental pedigree were performed, dividing it into two random halves with equal representation of parental treatment groups. Permutation revealed that 7.25% of enrichment for positive changes in genetic correlations in body size DE were attributed to pedigree structure rather than parental environment. It was significantly more likely that directional shifts in genetic correlation under parental upwelling were attributed to environment rather than pedigree (χ^2^ = 1639.6; *p* < 2e10^-16^). Most upwelling-induced directional shifts to genetic correlations in biomineralization DE (94.53%) were attributable to pedigree structure. Thus, there were detectable effects of parental environment on the genetic covariance of pathways associated with adaptive body size plasticity, but not biomineralization.

Heritable DE associated with plasticity in body size was enriched for functions associated with ribosomal biogenesis and maintenance (Fig. 6). Transcripts with high DE heritability and adaptive reductions in body size under upwelling were enriched for the BP/MF GO terms related to ribosomal biogenesis, RNA processing, and transmembrane transport localized to the nucleolus. Heritable DE associated with maladaptive increases in size was enriched for BP/MF terms related to amide formation (a component of translation) and ribosome structure localized to the cytosolic ribosome and large ribosomal subunit (Fig. 6). Heritable DE with adaptive effects on the plasticity of biomineralization was not enriched for GO terms.

**Figure 6.**
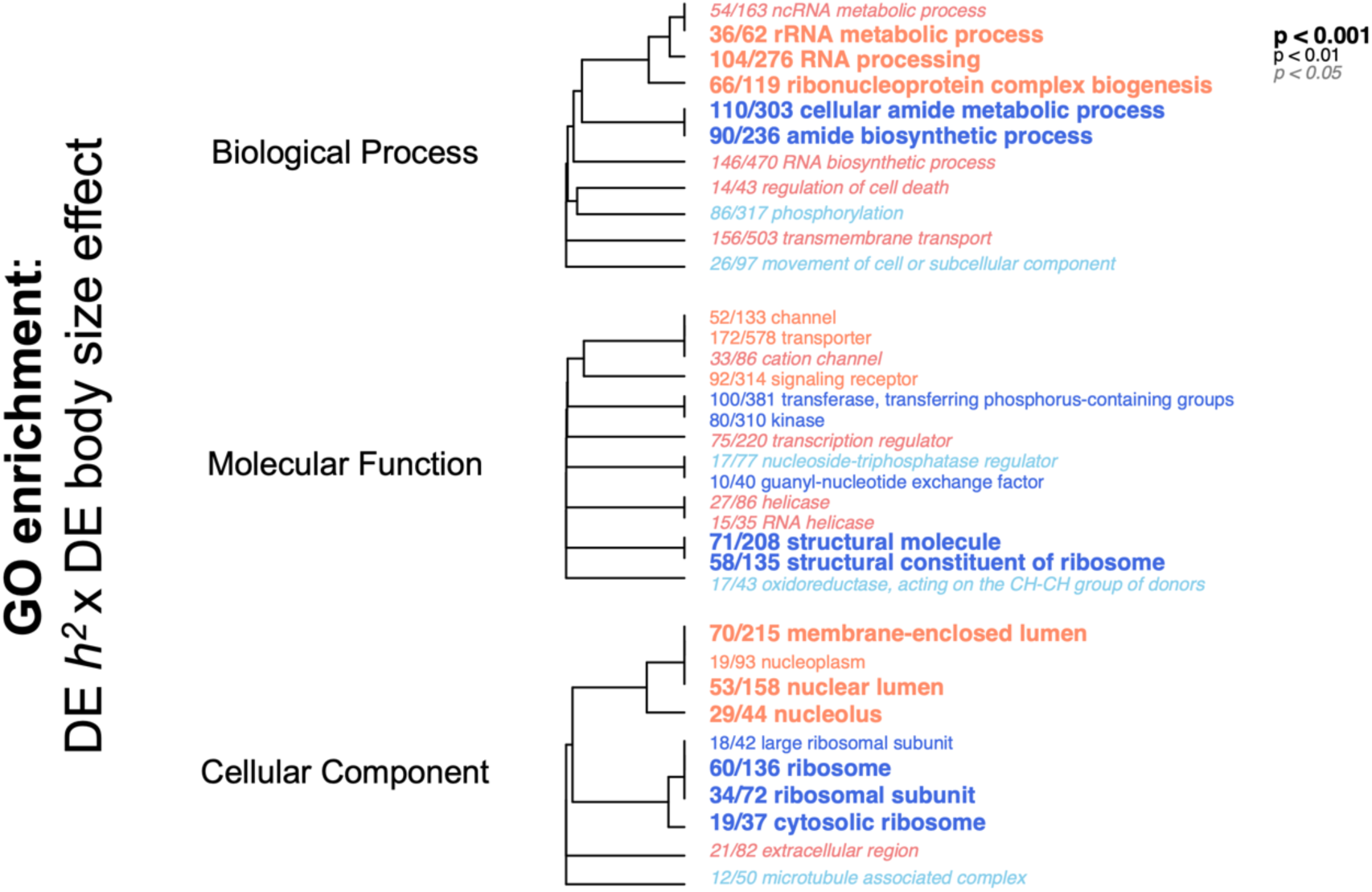
Functional enrichment of differentially expressed genes with high heritability and strong absolute effects on the plasticity of body size. Enriched gene ontology (GO) terms are derived from a Mann Whitney U test of variation in the product of differential expression’s heritability (h^2^) and DE’s effect on adaptive plasticity of body size in response to upwelling. Trees depict clustering of GO terms based on shared transcripts. Red GO terms are enriched within heritable differential expression associated with adaptive reductions in body size in response to developmental upwelling exposure. Blue terms are enriched within heritable differential expression associated with maladaptive upwelling exposure. Blue terms are enriched within heritable differential expression associated with maladaptive.

## 4. Discussion

### 4.1. Transgenerational effects reshaped fitness consequences of offspring performance traits and their plasticity

Conditioning of adult *S. purpuratus* to experimental upwelling during gametogenesis led to an increase in the strength of selection gradients acting on larval size, biomineralization, and the plasticity of both traits in response to larval upwelling exposure (Fig. 3). Transgenerational effects can influence the fitness consequences of offspring phenotypes through changes in provisioning to gametes (Gagliano and McCormick 2007). In these cases, it is expected that expression of a costly phenotype in offspring with fewer resources (e.g. protein, lipid, and amino acids) and lower condition will causes a fitness decline relative to the expression of that same phenotype in higher condition offspring (Roff and Fairbairn 2007). Maternal conditioning of *S. purpuratus* to ocean acidification has been shown to decrease lipid provisioning to eggs while leaving egg size and protein provisions unaffected (Wong et al. 2019). In our experiment, dams conditioned to different treatments did not did not differ in egg size (Strader et al. 2022). Our study showed that abnormal development (a proxy for larval mortality) was significantly higher in larvae from stress-exposed parents, demonstrating a clear decline in offspring condition (Fig. 3).

Our findings align with and expand on studies evaluating whether parental effects alter selection on offspring traits. Much of this literature has focused on parental care in nesting vertebrates that have substantial post-zygotic investment, while our study addressed prezygotic parental effects. In collared flycatchers, the strength and direction of selection acting on offspring size changed between low- and high-quality parental rearing conditions (Krist and Munclinger 2015). In blue tits, low parental provisioning toward hatched offspring strengthened negative selection on metabolic rate and begging behaviors (Lucass et al. 2016). In mice, maternal care weakened selection gradients acting on offspring traits by buffering against the effects of deleterious phenotypes (Ashbrook et al. 2015). An example of transgenerational effects on selection gradients driven by nutrient provisioning, rather than care, is in damselfish, where low maternal nutrition increased positive selection for increase yolk sac size in larval offspring (Gagliano and McCormick 2007). Our findings connect parental investment to selection on offspring traits through ecological physiology rather than parental care: parental exposure to abiotic stress reduced offspring condition, strengthening the fitness costs of multiple offspring traits that are critical for sustaining performance under stress.

Transgenerational plasticity (TGP) has been proposed to buffer against or facilitate natural selection in offspring phenotypes by masking or expressing genetic variation (Vogl 1996; Heckwolf et al. 2018). Our results demonstrate that transgenerational effects also reshape selection on offspring by increasing the fitness effects of expressed genetic variation. Expression of costly phenotypes induced by TGP in offspring that received poor provisioning may accelerate adaptation over multiple generations. While transgenerational effects have received interest in global change biology for their ability to promote rapid adaptive responses to climate stressors on ecological timescales, our findings align with recommendations that additional research should investigate their ability to modulate evolution and climate adaptation (Donelson et al. 2018; Fox et al. 2019). In populations of *S. purpuratus*, for example, local adaptation to ocean acidification is evident across a mosaic of upwelling in the California Current Ecosystem, where upwelling is predicted to become more frequent and severe under climate change (Snyder et al. 2003). Future research in natural populations of *S. purpuratus* assessing the effects of parental upwelling environment on adaptive evolution in larvae may reveal how transgenerational effects alter climate adaptation.

### 4.2. Transgenerational effects reshaped genetic variance-covariance in pathways associated with offspring performance

Genetic variance-covariance matrices (G matrices) can constrain responses to selection in multivariate trait space when genetic covariance or correlation between traits is strongly negative (Agrawal and Stinchcombe 2008). G matrices of differential expression associated with adaptive offspring plasticity shared a high degree of negative genetic correlation, demonstrating antagonistic pleiotropy or linkage between variants of opposing effects on gene expression traits (Chebib and Guillaume 2021). Pervasive negative genetic correlation indicates that adaptive responses to selection by some gene expression modules should be constrained (Sgrò and Hoffmann 2004). Among DE associated with adaptive plasticity in body size, genetic correlations between logFC values of thousands of genes significantly shifted from negative to positive in offspring of upwelling-conditioned parents (Fig. 5). Environments can influence genetic correlations between traits by inducing or silencing the expression of genetic variation underpinning those traits (Wood and Brodie 2015; Bogan and Yi 2024). For polygenic traits, this can lead to the expression of variants that share more positive or more negative genetic correlation. Effects of developmental environments on genetic correlation have been reported in pairs of traits (Sgrò and Hoffmann 2004; Wood and Brodie 2015; Fischer et al. 2016). Our results reveal that not only developmental, but parental environment, influences the expression of genetic covariation across thousands of transcriptional traits.

Increases in positive genetic correlation among DE associated with adaptive body size plasticity reduced genetic tradeoffs, mitigating constraints on adaptive evolution (Fig. 5). Shifts toward more positive genetic correlations in trait-associated pathways in larvae from upwelling-exposed parents coincided with increases in the strength of selection acting on those traits. This combination of increased selective pressure and reduced constraints on responses to selection suggest that parental upwelling exposure acted as a facilitator of adaptive evolution. Models of adaptation to environmental change frequently fit an effect of each generation’s environment on the expression of genetic (co)variation and selection gradients (Gillberg et al. 2019; Cubry et al. 2022; Gallegos et al. 2025). However, the influence of environmental experience in prior generations on these parameters in descendent generations is rarely considered. Our results motivate consideration of parental environments and their transgenerational effects as processes reshaping genetic evolution and adaptation in later generations, particularly in cases of adaptation to global change.

### 4.3. Reduced heritability of transcriptional responses associated with adaptive plasticity

Parental effects on the selection gradients and genetic covariance of offspring performance traits will only influence evolution if there is heritable genetic variation underpinning those traits. We observed that the heritability of DE significantly decreased when it was associated with adaptive, phenotypic responses to larval upwelling conditioning (Fig. 4). This extent of genetic variation for DE, and its decline in DE associated with adaptive plasticity, confirms observations in other studies of plants (He et al. 2021) and animals (Oostra et al. 2018; Campbell-Staton et al. 2021). However, past research has assigned putatively adaptive and maladaptive roles to molecular responses based on prior findings. Here, we directly integrated gene expression plasticity with fitness measures. Quantifying cross-environment genetic correlations (a measure of genetic variance in plasticity) for gene expression in a tropical butterfly, Oostra et al. detected low-to-undetectable genetic variation for DE between two seasonal morphological phenotypes (Oostra et al. 2018). The authors proposed that minimal genetic variation for transcriptional plasticity resulted from a canalized response to a highly reliable seasonal cue (Oostra et al. 2018). Numerous studies have reported moderate-to-high genetic variation in DE measured as genotype x environment (GxE) interactions shaping expression in animals (Grishkevich and Yanai 2013; McCairns et al. 2016; Huang et al. 2020). Oostra et al. provide one of the only measures of additive genetic variance in the reaction norm of gene expression rather than variance attributed to GxE. Thus, it is notable that V_A_ of DE that they detected is strikingly lower than what we observed here.

*S. purpuratus* exhibited reduced genetic variation for transcriptional plasticity associated with adaptive phenotypic effects, but heritability remained significant for a proportion of these genes (Fig. 4). This pattern is potentially a signature of standing genetic variation and selection for DE induced by upwelling in natural populations. *S. purpuratus* larvae are widely dispersed during their planktonic phase resulting in high connectivity across spatial scales and high genetic diversity within populations (Palumbi and Wilson 1990; Edmands et al. 1996; Pespeni and Palumbi 2013). Its dispersal distances can be wide enough that larvae are frequently transported across areas of major and minor upwelling in the California Current and Southern California Bight (Zaytsev et al. 2003; Pespeni and Palumbi 2013; Chan et al. 2017). *S. purpuratus* populations exhibit evidence of local adaptation to regional differences in *p*CO_2_ despite high rates of gene flow (Evans et al. 2013; Pespeni et al. 2013). Balancing selection during seasonal changes in upwelling intensity may also sustain genetic variation in plasticity (Lynn and Simpson 1987; Quilfen et al. 2021). Prior research on *S. purpuratus* suggests that an individual population should possess genetic variation in plastic responses associated with coastal upwelling attributed to migration and seasonal variation in selection, but that reductions in genetic variation for adaptive plasticity may occur due to selection.

DE that was heritable and associated with adaptive plasticity, and whose genetic correlations were made more positive by parental upwelling, were enriched for specific functions. Heritable and adaptive changes in gene expression were enriched for GO terms associated with ribosomal biogenesis while maladaptive, heritable DEGs retained enriched functions indicative of ribotoxic stress such as downregulated ribosomal subunits (Fig. 6). Induced ribotoxic stress can result in the downregulation of ribosomal subunits and the activation of HSF1, which upregulates the cytosolic, 70 kda heat shock proteins Hsc70 and Hsp40 (Albert et al. 2019). These are the two classes of heat shock proteins that were upregulated in response to larval upwelling conditioning. Abiotic stress can perturbate ribosomal function via denaturation of ribosomal proteins/RNAs or misfolding of nascent proteins that disrupt proximal ribosomes (Iordanov et al. 1998; De and Mühlemann 2022). The overrepresentation of genes involved in ribosomal biogenesis and structure among transcripts with heritable, (mal)adaptive plasticity in larval body size may be explained by essential role of ribosomal homeostasis for growth and development (MacInnes 2016). Perturbations in ribosomal function during development bear harmful effects on organismal function and size (Ordas et al. 2008; Freed et al. 2010).

## 5. Conclusion

Understanding how environments reshape the expression of genetic variation is a major frontier in biology (Gibson and Dworkin 2004; Riederer et al. 2022), one that can improve forecasts of adaptation (Martin et al. 2023). Prior research has focused on developmental environments and within-generation processes, while less is understood about transgenerational effects (Goldstein and Ehrenreich 2021). We demonstrated that parental environments not only influence the expression of genetic variation in offspring traits, but the strength of selection on those traits as well. Parental exposure to upwelling in *S. purpuratus* (i) increased the strength of selection on offspring performance traits and their plasticity in response to developmental upwelling exposure and (ii) caused negative-to-positive shifts in genetic covariance between gene expression traits associated with adaptive plasticity. Combined, increases in the strength of selection and reductions in genetic tradeoffs between adaptive traits are expected to accelerate rates of adaptive evolution (Agrawal and Stinchcombe 2008; Blows and McGuigan 2016). Coastal upwelling has shaped local adaptation among *S. purpuratus* populations in the California Current Ecosystem (Kelly et al. 2013; Pespeni and Palumbi 2013; Evans et al. 2017), where upwelling is predicted to become more severe under climate change (Quilfen et al. 2021). Our results suggest that transgenerational effects on selection and genetic (co)variation potentially shaped adaptation to upwelling in *S. purpuratus* and may alter the course of evolution under changing upwelling regimes. More broadly, our experiment suggests that parental environments and transgenerational effects play significant roles in mediating evolution by natural selection.

## Acknowledgements

We thank Juliet Wong, Logan Kozal, Terence Leach, Jannine Chamorro, Maddie Housh, and Olivia Simon for their assistance in adult conditioning and experimental culturing. We thank Matthew Wolak, Katie Lotterhos, Steven Roberts, Yaamini Venkataraman, and Alan Downey-Wall for his insights on quantitative genetic and transcriptomic analyses. This research was funded by United States National Science Foundation awards IOS-1656262 and OPP-2053726 to GEH. Diving and boating resources were provided by Santa Barbara Coastal Long Term Ecological Research program (NSF award OCE-1831937; Director: Dr. Robert Miller).

## Data availability

Raw RNA-seq data are available on the Short Read Archive under Bioproject <u>PRJNA1169019</u>. All phenotypic data, pedigree data, and code are hosted on the Github repository https://github.com/snbogan/QG_Purp_RNA. An archived copy of this Github repository is available on Zenodo under the DOI [will be produced upon acceptance].

## Author Contributions

SNB, MS, and GEH conceived of the aims and scope of the study. MS led experimental design and phenotyping. GEH assisted SB assisted with experimental culturing. SB and MS assisted with seawater chemistry and phenotyping. SB performed RNA extractions, oversaw sequencing, bioinformatics, and all statistical analyses. SB wrote the manuscript. All authors edited and revised the manuscript.

## Declarations

The authors declare no competing interests.

