## Supplemental Materials for "Parental stress alters the fitness effects and genetic correlation of offspring performance traits and their gene expression pathways"

**Running head:** Parental stress shapes adaptive evolution

*Supplemental Methods*

*1.1. RNA-seq processing and alignment* - Illumina Universal Adapters were removed from paired end reads using CutAdapt v4.4 (Martin 2011) and reads were trimmed and quality filtered using Trimmomatic v0.39 set to paired end mode, trailing and leading clip = 3, sliding window = 4:15, minimum length = 36, and headcrop = 10 (Bolger et al. 2014). All trimmed reads passed quality check via FastQC v0.12.1 (Andrews). Forward strands of paired reads were aligned to the 'Spur_5.1' reference genome assembly (Sodergren et al. 2006) using hisat2 v2.2.1 (Kim et al. 2019). Forward rather than paired reads were aligned to increase the number of genes included in the dataset after filtering for read count. Resulting SAM alignments were sorted and converted to BAM using SAMtools v1.6 (Li et al. 2009). Reads were counted per transcript from sorted BAM files using featureCounts v1.6.3 input with the 'Spur_5.1' gtf annotation set to a MAPQ alignment quality cutoff of 10 (Liao et al. 2014). Read counts were performed with transcript annotations rather than genes to ensure accurate estimation of additive genetic variation for transcriptional traits, which may vary between transcript isoforms. Transcripts of the same parent gene made up 9.14% of the final filtered data set used for differential expression analysis and quantitative genetic models. DE transcripts associated with adaptive plasticity in larval body size and biomineralization included 0 pairs of transcript isoforms of the same parent gene, avoiding any confounds of genetic correlation.

*Supplemental Results*

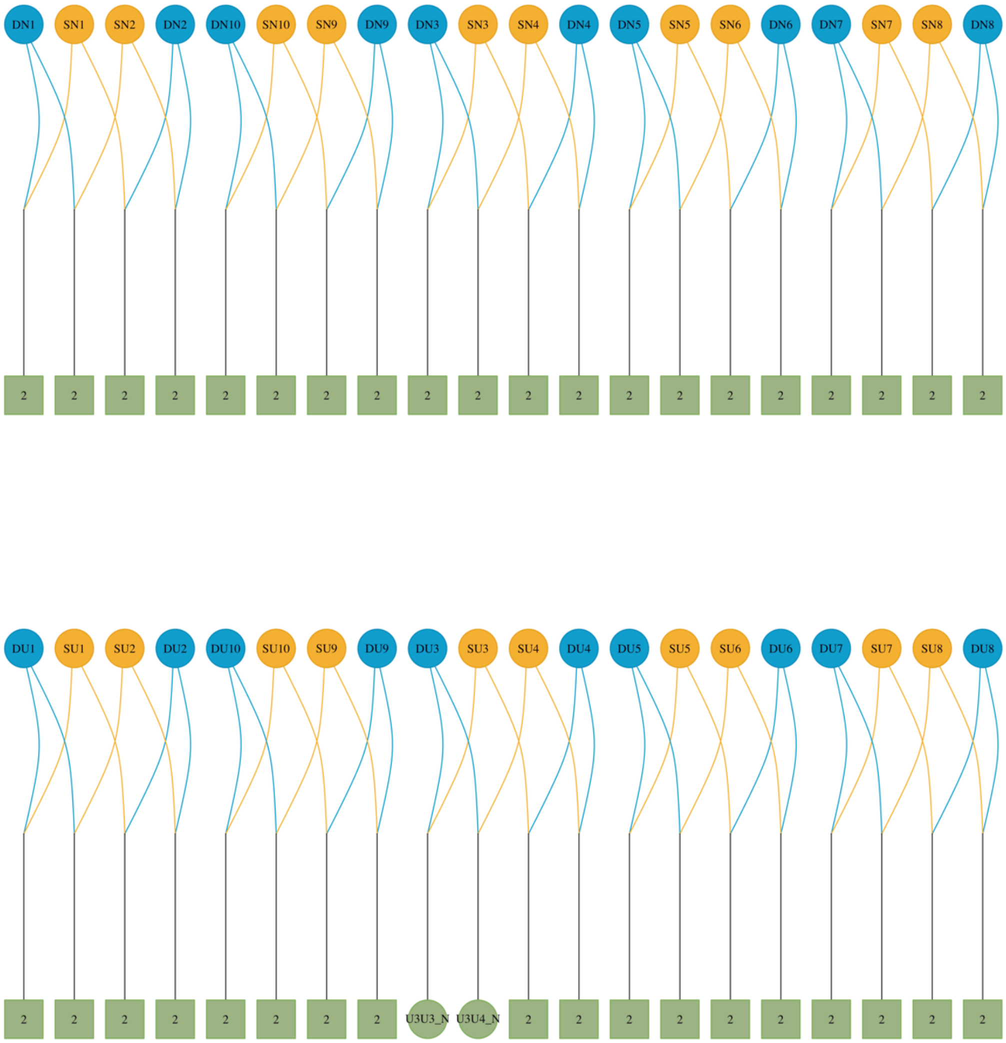


**Figure S1** | *Pedigree and experimental breeding design.* The pedigree is split between families spawned from non-upwelling parents (top) and upwelling parents (bottom). Dams are colored in blue. Sires are colored in orange. Breeding is represented by the connection of colored lines between dams and sires. Black lines depict larval offspring. Larval families that were split and conditioned to both upwelling and non-upwelling developmental conditions are represented by boxes labeled with "2". Families for which a developmental treatment group was filtered out during RNA-seq outlier removal are labeled as circles and the retained replicate ID. Dams and sires are respectively labeled with "D" and "S" followed by "N" (parental non-upwelling) or "U" (parental upwelling). Larval families are labeled first by dam ID, then sire ID, followed by their developmental treatment.

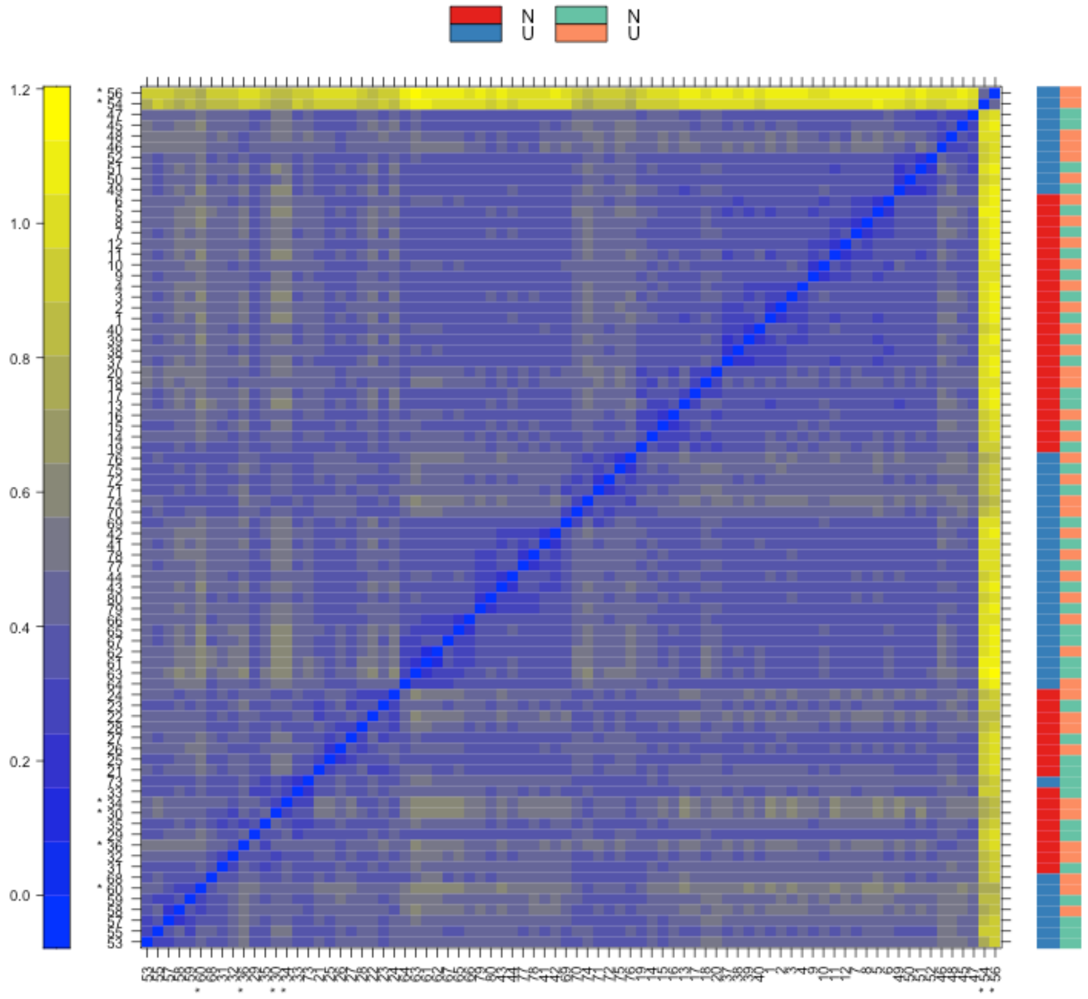


**Figure S2** | *Gene expression distances between samples calculated by arrayQualityMetrics.* A false color heatmap of the distances between arrays. Parental environment is represented by blue for non-upwelling (N) and red for upwelling (U). Developmental environments are represented by N = green and U = pink. The color scale is chosen to cover the range of distances encountered in the dataset. Patterns in this plot can indicate clustering of the arrays either because of intended biological or unintended experimental factors (batch effects). The distance $d_{ab}$ between two arrays $a$ and $b$ is computed as the mean absolute difference ($L_1$-distance) between the data of the arrays (using the data from all probes without filtering). In formula, $d_{ab}$ = mean | $M_{ai}$ - $M_{bi}$ |, where $M_{ai}$ is the value of the $i$-th probe on the $a$-th array. Outlier detection was performed by looking for arrays for which the sum of the distances to all other arrays, $S_a = \Sigma_b d_{ab}$ was exceptionally large. 6 such arrays were detected, and they are marked by an asterisk, *.

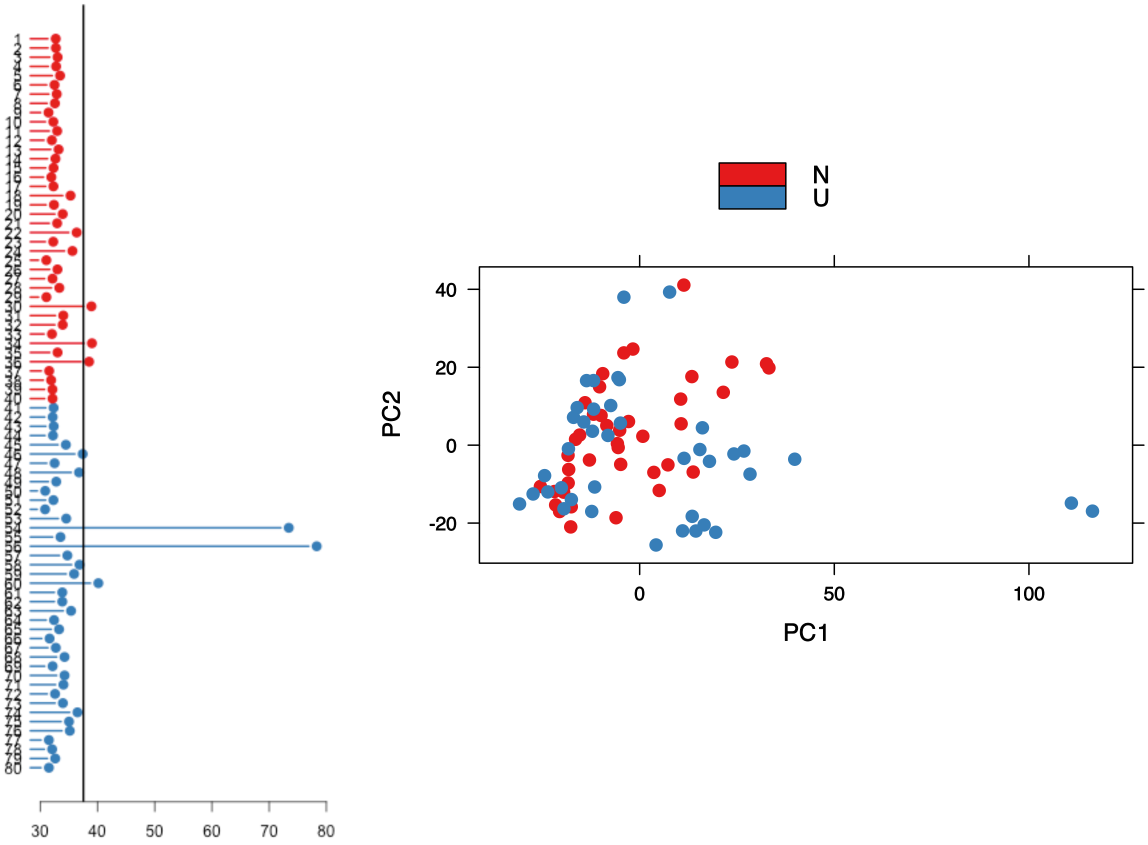


**Figure S3 |** *Outlier detection derived from gene expression distances and principal components analysis performed in arrayQualityMetrics*. (Left) A bar chart of the sum of distances to other arrays $S_a$, the outlier detection criterion from the previous figure. The bars are shown in the original order of the arrays. Based on the distribution of the values across all arrays, a threshold of 37.5 was determined, which is indicated by the vertical line. 6 arrays exceeded the threshold and were considered outliers. (Right) A scatterplot of the arrays along the first two principal components. You can use this plot to explore if the arrays cluster, and whether this is according to an intended experimental factor, or according to unintended causes such as batch effects. Move the mouse over the points to see the sample names. Principal component analysis is a dimension reduction and visualization technique that is here used to project the multivariate data vector of each array into a two-dimensional plot, such that the spatial arrangement of the points in the plot reflects the overall data (dis)similarity between the arrays.

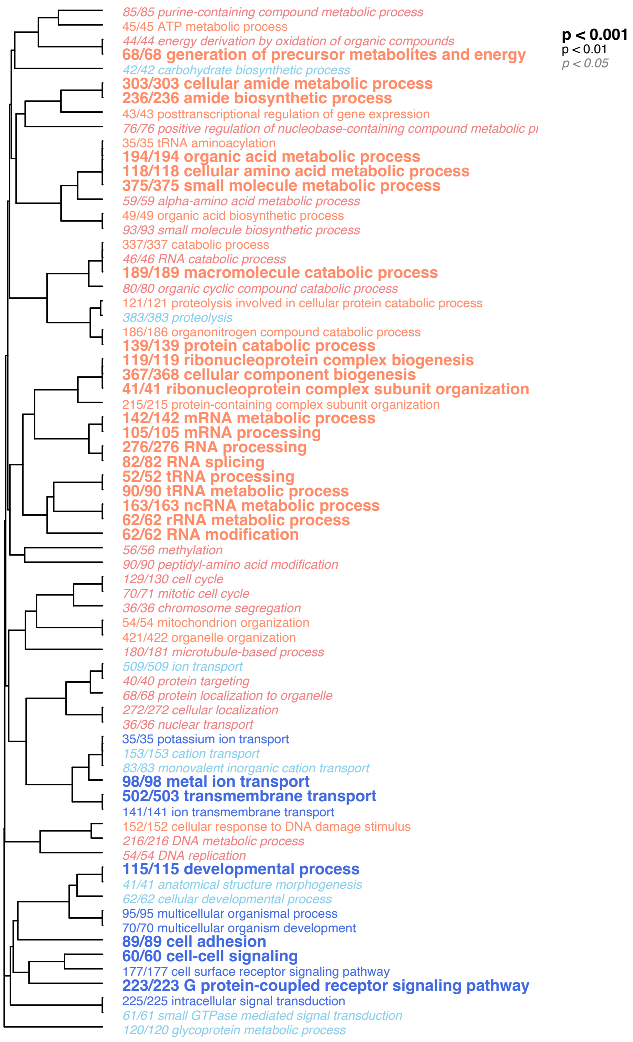


**Figure S4** | *Enriched biological process (BP) gene ontology (GO) among transcripts differentially expressed in response to parental upwelling conditions*. BP GO terms are clustered according to the number of transcripts common to each enriched term. P-values are associated with GO term enrichment among upregulation (red) or downregulation (blue).

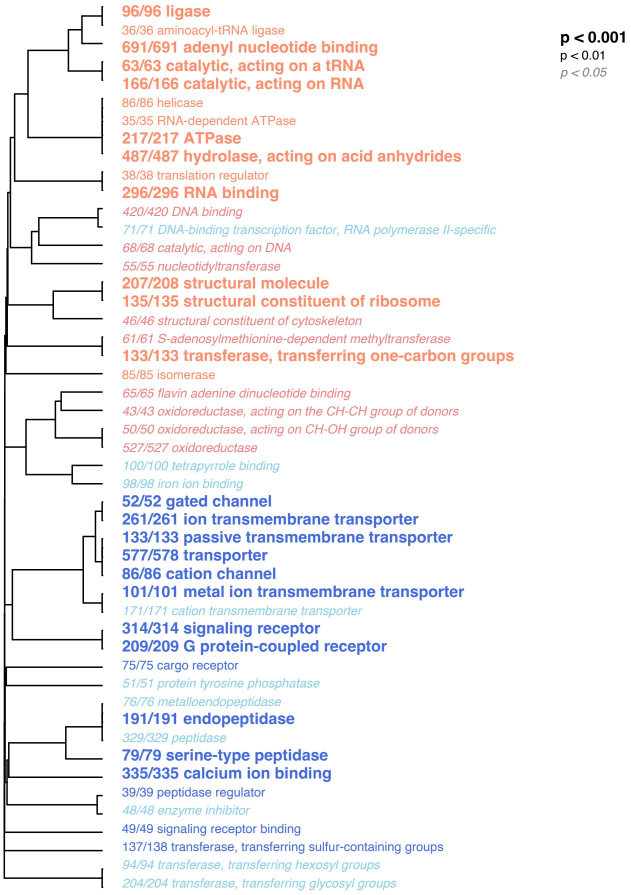


**Figure S5** | *Enriched molecular function (MF) gene ontology (GO) among transcripts differentially expressed in response to parental upwelling conditions*. MF GO terms are clustered according to the number of transcripts common to each enriched term. P-values are associated with GO term enrichment among upregulation (red) or downregulation (blue).

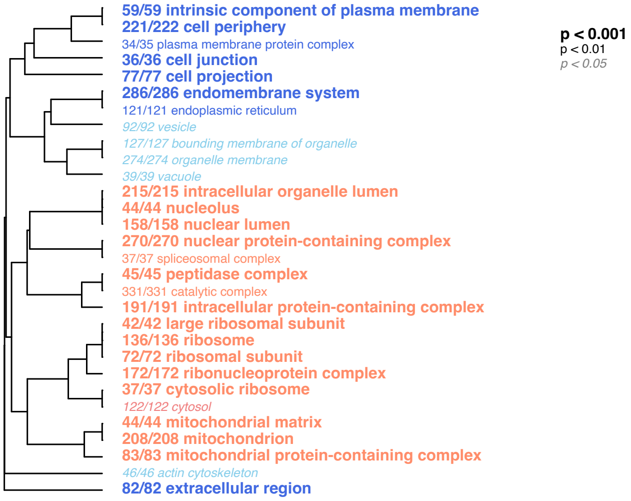


**Figure S6** | *Enriched cellular component (CC) gene ontology (GO) among transcripts differentially expressed in response to parental upwelling conditions*. CC GO terms are clustered according to the number of transcripts common to each enriched term. P-values are associated with GO term enrichment among upregulation (red) or downregulation (blue).

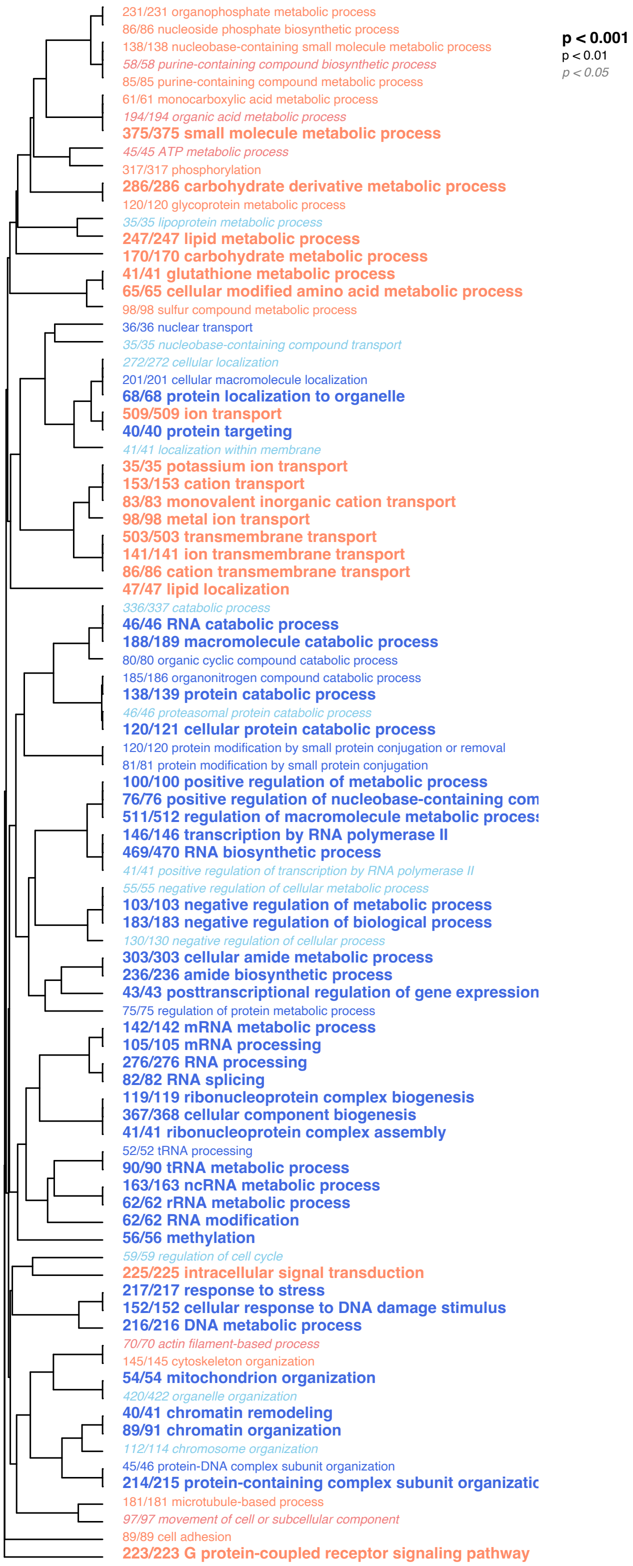


**Figure S7** | *Enriched biological process (BP) gene ontology (GO) among transcripts differentially expressed in response to developmental upwelling conditions*. BP GO terms are clustered according to the number of transcripts common to each enriched term. P-values are associated with GO term enrichment among upregulation (red) or downregulation (blue).

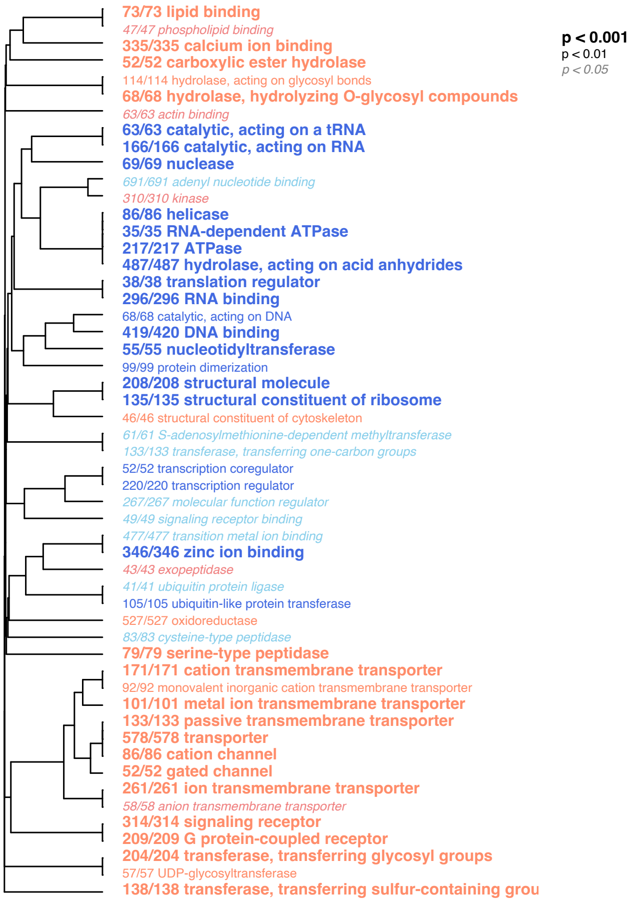


**Figure S8** | *Enriched molecular function (MF) gene ontology (GO) among transcripts differentially expressed in response to developmental upwelling conditions*. MF GO terms are clustered according to the number of transcripts common to each enriched term. P-values are associated with GO term enrichment among upregulation (red) or downregulation (blue).

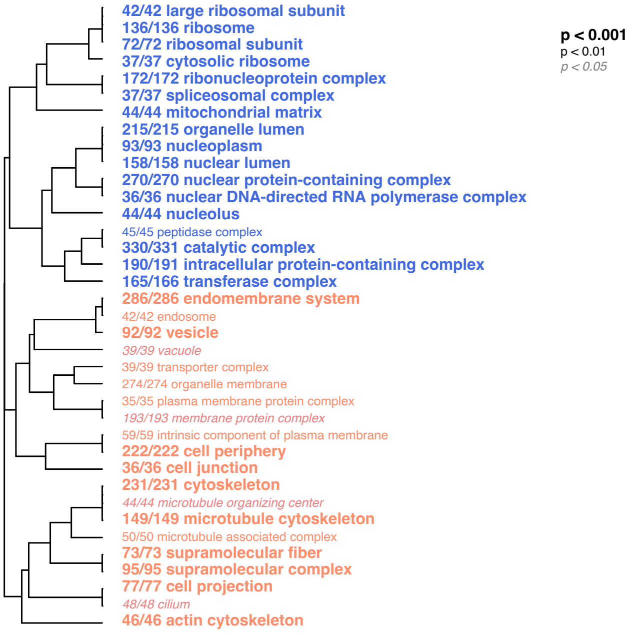


**Figure S9** | *Enriched cellular component (CC) gene ontology (GO) among transcripts differentially expressed in response to developmental upwelling conditions*. CC GO terms are clustered according to the number of transcripts common to each enriched term. P-values are associated with GO term enrichment among upregulation (red) or downregulation (blue).
